# Circadian Rhythm Is Disrupted by ZNF704 in Breast Carcinogenesis

**DOI:** 10.1101/2020.01.09.900548

**Authors:** Chao Yang, Jiajing Wu, Xinhua Liu, Yue Wang, Beibei Liu, Xing Chen, Xiaodi Wu, Dong Yan, Lulu Han, Shumeng Liu, Lin Shan, Yongfeng Shang

## Abstract

Copy number gain in chromosome 8q21 is considered as the prototype of genetic abnormalities associated with development of breast cancer, yet the oncogenic potential underlying this amplicon in breast carcinogenesis remains to be delineated. We report here that *ZNF704*, a gene mapped to 8q21, is recurrently amplified in various malignancies including breast cancer. We found that ZNF704 acts as transcription repressor and interacts with the transcription corepressor SIN3A complex. Genome-wide interrogation of the transcriptional targets identifies that the ZNF704/SIN3A complex represses a panel of genes including *PER2* that are critically involved in the function of circadian clock. Indeed, ZNF704 overexpression prolongs the period and dampens the amplitude of circadian clock. We showed that ZNF704 promotes the proliferation and invasion of breast cancer cells *in vitro* and accelerates the growth and metastasis of breast cancer *in vivo*. Consistently, the level of ZNF704 expression is inversely correlated with that of PER2 in breast carcinomas, and high level of ZNF704 correlates with advanced histological grades, lymph node positivity, and poor prognosis of breast cancer patients, especially those with HER2^+^ and basal-like subtypes. These results indicate that ZNF704 is an important regulator of circadian clock and a potential driver for breast carcinogenesis.

## Introduction

Structural and numerical alterations of chromosome 8 have been reported in up to 60% of breast cancer cases(Courjal & Theillet, 1997; Tirkkonen et al, 1998), and copy number gains involving the long arm of chromosome 8, including high-level amplifications at 8q21 and 8q24, are considered to be the prototype of genetic abnormalities associated with development of breast cancer as well as cancers from other tissue origins and also with poor prognosis of patients(Balleine et al, 2000; Choschzick et al, 2010; Raeder et al, 2013). While the role of the MYC gene as the driver of the 8q24 amplicon is well established, the genetic factor(s) contributing to the oncogenic potential of the 8q21 amplicon remains to be elucidated. It is reported that amplification of the gene encoding for WW domain-containing E3 ubiquitin protein ligase 1 (WWP1) in this region is an oncogenic factor for breast cancer(Chen et al, 2010) and prostate cancer(Chen et al, 2007), while amplification of the gene encoding for tumor protein D52 (TPD52), whose function has rarely been studied, in 8q21 is implicated in the development of ovarian cancer(Byrne et al, 2010) and lung cancer(Kumamoto et al, 2016). Clearly, the molecular basis underlying the 8q21 amplicon in the development and progression of breast carcinogenesis needs further elucidation.

Circadian rhythm is generated via oscillations in the expression of clock genes that are organized into a complex transcriptional-translational autoregulatory network to dictate an array of physiological and behavioral activities in responding to periodic environmental changes(Bass & Takahashi, 2010; Takahashi, 2016). Central to the molecular system controlling the circadian rhythm is the heterodimer of transcription factors, BMAL1 (Brain and Muscle ARNT-Like 1, also known as ARNTL) and CLOCK (the circadian locomotor output cycles kaput), which activates the transcription of genes containing E-box binding sequences in their promoter/enhancer regions, including *Period* (*PER1*, *PER2*) and *Cryptochrome* (*CRY1*, *CRY2*), and PER1/2 and CRY1/2, in turn, heterodimerizes with BMAL1/CLOCK to inhibit their own transcription(Hida et al, 2000; Kume et al, 1999; Reppert & Weaver, 2002). Given the paramount importance of circadian clock in the regulation of cellular activities and in the maintenance of cell homeostasis, its contribution to the pathogenesis of several diseases is highly predicted. Indeed, animal models and epidemiological studies suggest that dysfunction of circadian clock is associated with increased incidences of various epithelial cancers(Filipski et al, 2002; Schernhammer et al, 2001; Viswanathan et al, 2007), and aberrant expression of core clock genes is found in a broad spectrum of malignancies including breast cancer(Chen et al, 2005), glioma(Luo et al, 2012), leukemia(Taniguchi et al, 2009), and colorectal cancer(Yu et al, 2013). Clearly, understanding the regulation/deregulation of clock gene expression is of great importance to the understanding of the molecular carcinogenesis.

*PER2* is an indispensable clock gene that constitutes the negative limb in the transcriptional-translational feedback loop of the circadian clock(Bae et al, 2001; Dunlap, 1999). Interestingly, *PER2* plays an important role in the control of cellular proliferation and has been suggested to be a tumor suppressor(Fu et al, 2002). *PER2* expression is significantly reduced in both sporadic and familial primary breast cancers(Winter et al, 2007), and deficiency of *PER2* affects the growth rate in silkworm(Sandrelli et al, 2007) and accelerates the proliferation of breast cancer cells and the growth of breast cancer by altering the daily growth rhythm(Yang et al, 2009). At the cellular level, *PER2* controls lipid metabolism and adipocyte cell differentiation through direct regulation of PPARγ; lack of PER2 leads to the cellular differentiation from fibroblast to adipocyte(Grimaldi et al, 2010). At the molecular level, PER2 was shown to repress the transcription of *TWIST* and *SLUG* to inhibit epithelial-mesenchymal transition(Hwang-Verslues et al, 2013), a key step leading to cancer metastasis(Wang & Shang, 2013). Thus, understanding the regulation/deregulation of PER2 expression is important to the understanding of its role in tumorigenesis.

In this study, we investigated the oncogenic potential of the 8q21 amplicon. We found that *ZNF704*, a gene that is mapped to 8q21, is frequently amplified in various cancers. We showed that at the molecular level ZNF704 acts in concert with the SIN3A complex to repress the transcription of PER2, an essential component of the molecular system that controls circadian rhythm. We demonstrated that ZNF704 disrupts the circadian rhythm and promotes breast carcinogenesis.

## Results

### *ZNF704*, a Gene Harbored in the 8q21 Amplicon, Is Amplified/Overexpressed in a Variety of Cancers

As stated above, although amplification of chromosome 8q21 is a frequent event in various of cancers and is associated with poor prognosis of patients(Balleine et al, 2000; Choschzick et al, 2010), the genetic factor(s) that contribute to its oncogenic potential remain to be delineated. As the epigenetic mechanisms underlying the transcription regulation and the molecular basis underlying breast carcinogenesis are the primary focuses of our laboratory(Shan et al, 2016; Si et al, 2015; Wang et al, 2009; Yan et al, 2015; Zhang et al, 2016), we noted that one gene in the 8q21 region, *ZNF704*, which encodes for a zinc finger transcription factor, exhibited various genetic abnormalities in a broad spectrum of malignancies including cancers originated from prostate, liver, breast, uterus, and lung (Figure 1A), as bioinformatics analysis of the public datasets in the cBioPortal for Cancer Genomics (http://www.cbioportal.org/) indicated. Notably, amplification of *ZNF704* is the most frequent event across the abnormalities in the majority of the cancer types, occurring in ∼8% cases in prostate cancer, liver cancer, and breast cancer (Figure 1A). In concordance, analysis of the public datasets in Oncomine (https://www.oncomine.org/) showed that ZNF704 is significantly overexpressed in breast, liver, and prostate cancer (Figure 1B). Further analysis of two public datasets(2012; Ciriello et al, 2015) from cBioPortal for Cancer Genomics indicates that amplification of chromosome 8q21 region in breast carcinomas encompasses *ZNF704* loci (Figure 1C), and analysis of the public datasets (GSE9014, GSE72653, and GSE27567) showed that ZNF704 is upregulated in breast cancer samples (Figure 1D). Together, these observations support a notion that *ZNF704* is amplified/overexpressed in breast cancer.

**Figure 1.**
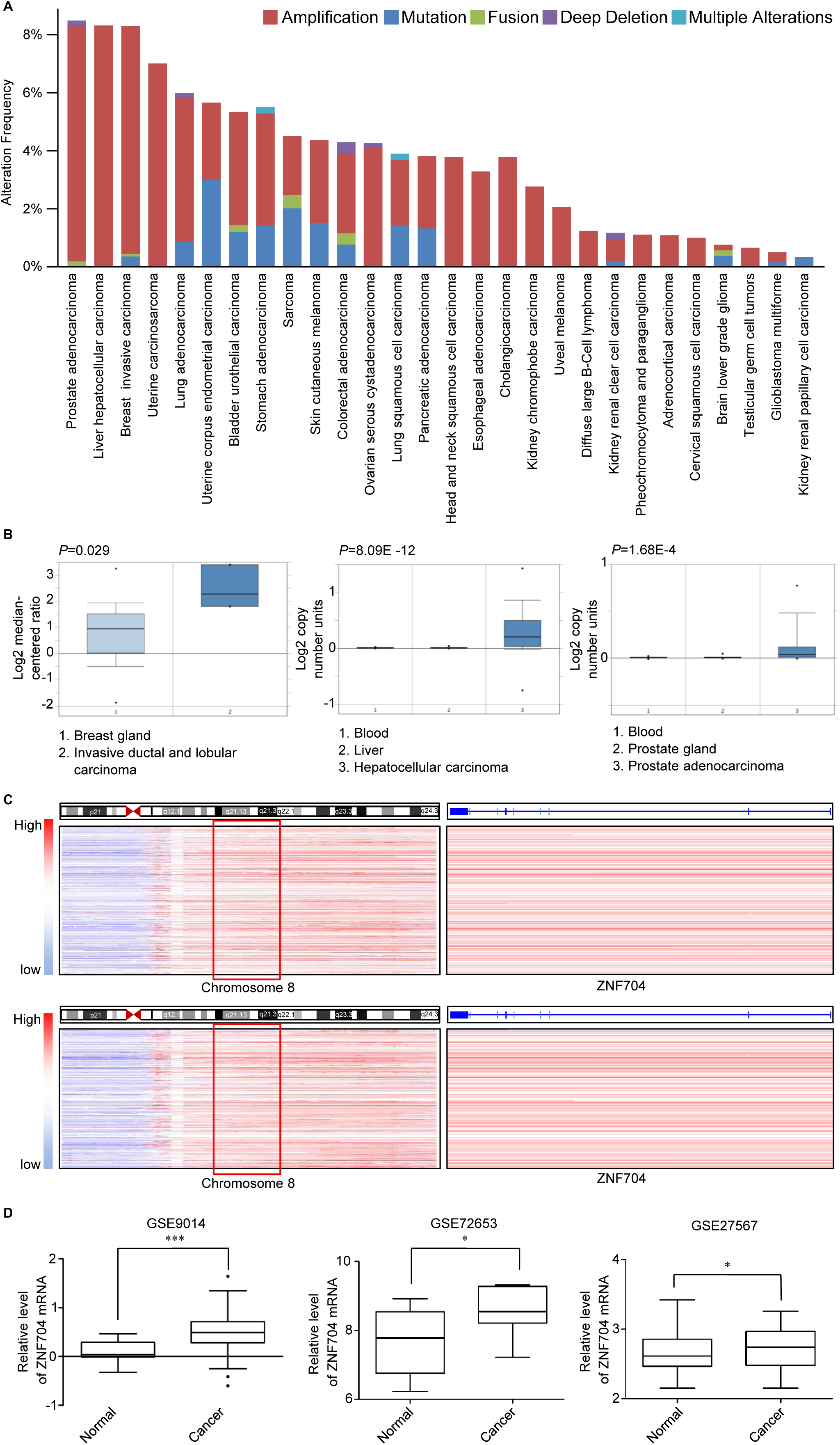
ZNF704 Is Amplified/Overexpressed in a Variety of Cancers. (A) Analysis of genetic alterations of *ZNF704* in a series of cancers from cBioPortal for Cancer Genomics (http://www.cbioportal.org/). (B) Analysis of TCGA datasets in Oncomine (https://www.oncomine.org/) for the expression or copy number of *ZNF704* between tumor and normal tissues. (C) Analysis of two public datasets from cBioPortal for Cancer Genomics in 2015 (upper) and 2012 (lower) for the amplification of 8q21 region and *ZNF704* in breast cancer patients. (D) Bioinformatics analysis of the public datasets (GSE9014, GSE72653 and GSE27567) in breast carcinoma samples and normal tissues. Data information: In (D), data are presented as mean ± SEM. \**P* <0.05, \*\*\**P* < 0.001 (Student’s t-test).

### ZNF704 Is a Transcription Repressor and Physically Associated with the SIN3A Complex

To explore the cellular function of ZNF704, we first cloned the gene encoding for human ZNF704 from a human mammary cDNA library (Clontech). To confirm the expression of ZNF704 protein, FLAG-tagged ZNF704 expression plasmid (FLAG-ZNF704) was transfected into MCF-7 or HEK293T cells. Cellular proteins were extracted from these cells as well as from several other cell lines and analyzed by western blotting with a monoclonal antibody against FLAG or polyclonal antibodies against ZNF704. The results showed that endogenous ZNF704 is a protein with a molecular weight of ∼60 kDa (Figure 2A), and that ZNF704 is expressed at variable levels in different cell lines (Figure 2B). Immunofluorescent imaging of ZNF704 in MCF-7 cells indicates that ZNF704 is primarily localized in the nucleus (Figure 2C).

**Figure 2.**
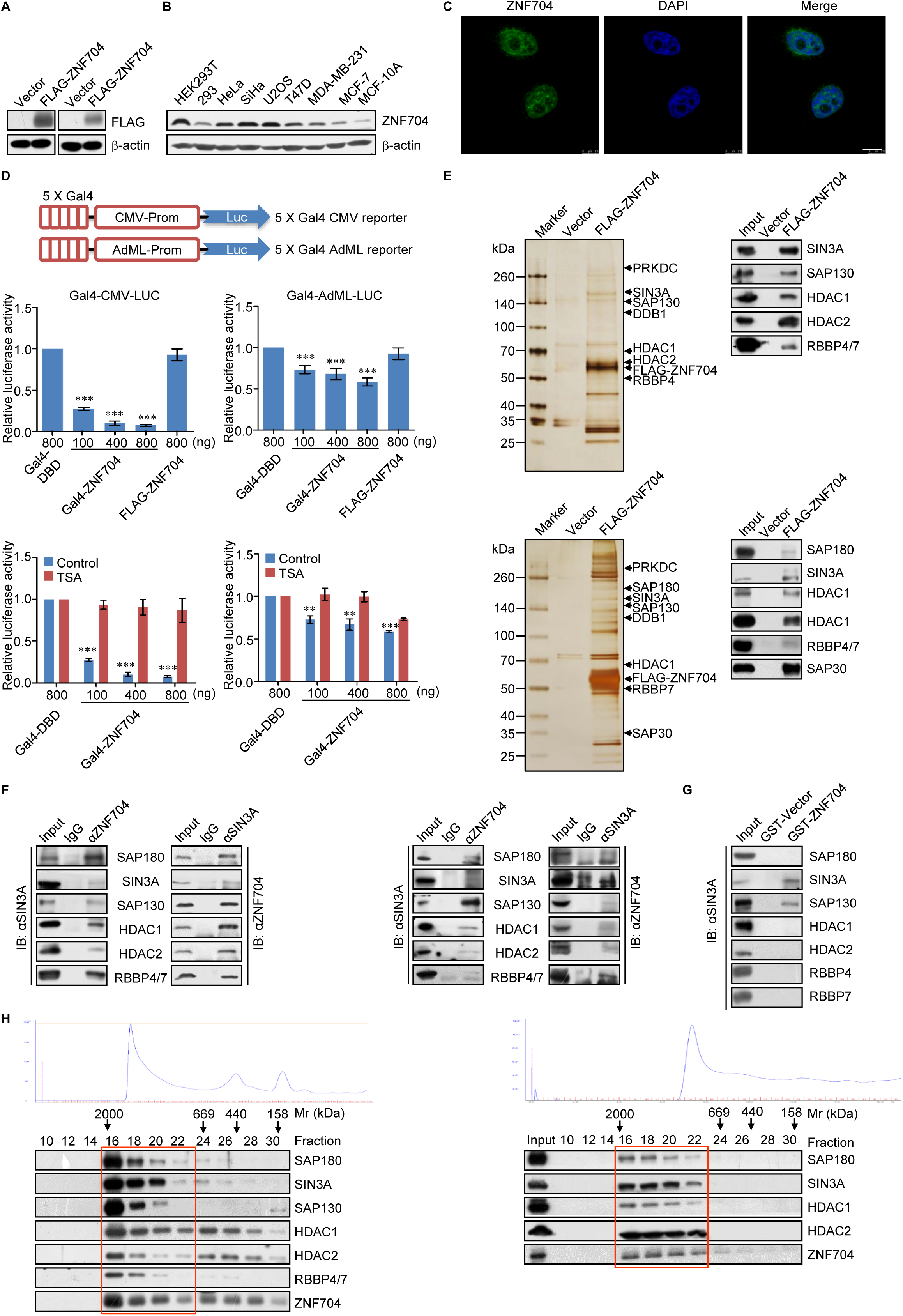
ZNF704 Is a Transcription Repressor and Physically Associated with the SIN3A Complex. (A) HEK293T (left) or MCF-7 (right) cells were transfected with empty vector or FLAG-ZNF704 for western blotting with antibodies against FLAG or β-actin. (B) Cellular proteins were extracted from the indicated cell lines for western blotting with antibodies against ZNF704 or β-actin. (C) The distribution of endogenous ZNF704 was detected by immunofluorescent microscopy. Bar: 7.5 μm. (D) Schematic diagrams of the Gal4-luciferase reporter constructs (upper). For reporter assays, HeLa cells were transfected with different amounts of Gal4-ZNF704 or FLAG-ZNF704 together with the indicated Gal4-luciferase reporter with or without treatment of TSA (lower). Each bar represents mean ± SD for triplicate experiments (\*\**P* < 0.01, \*\*\**P* < 0.001). (E) Immunopurification and mass spectrometric analysis of ZNF704-associated proteins in MDA-MB-231 (upper) and HEK293T (lower) cells. Cellular extracts from MDA-MB-231 or HEK293T cells stably expressing FLAG-ZNF704 were subjected to affinity purification with anti-FLAG affinity columns and eluted with FLAG peptides. The eluates were resolved by SDS-PAGE and silver stained. The protein bands were retrieved and analyzed by mass spectrometry (left); Column-bound proteins were analyzed by western blotting using antibodies against the indicated proteins (right). (F) Co-immunoprecipitation in MDA-MB-231 (left) and MCF-7 (right) cells with anti-ZNF704 followed by immunoblotting with antibodies against the indicated proteins, or immunoprecipitation with antibodies against the indicated proteins followed by immunoblotting with antibodies against ZNF704. (G) GST pull-down assays with GST-fused ZNF704 and *in vitro* transcribed/translated proteins as indicated. (H) FPLC analysis of nuclear extracts from MDA-MB-231 cells (left) and analysis of FLAG-ZNF704 affinity eluates in MDA-MB-231 cells stably expressing FLAG-ZNF704 (right). Chromatographic elution profiles and immunoblotting analysis of the chromatographic fractions are shown. Equal volume from each fraction was analyzed, and the elution position of calibration proteins with known molecular masses (kilodaltons) are indicated. Western blotting of ZNF704-containing complex fractionated by Superose 6 gel filtration.

We next determined the transcriptional activity of ZNF704. For this purpose, full-length ZNF704 was fused to the C terminus of the Gal4 DNA-binding domain (Gal4-ZNF704), and the transcriptional activity of the fused construct was tested in HeLa cells. We used two different Gal4-driven luciferase reporter systems, both contain 5 copies of the Gal4 binding sequence but differ in basal promoter elements (Figure 2D, upper). The results showed that Gal4-ZNF704 elicited a robust repression of the reporter activity in a dose-dependent fashion in both of the reporter systems, whereas overexpression of FLAG-ZNF704 had no effect on the activity of the Gal4-driven reporters (Figure 2D, lower), suggesting that ZNF704 must be physically associated with DNA to exert its transcription repression activity. In addition, treatment of HeLa cells with trichostatin A (TSA), a specific histone deacetylase (HDAC) inhibitor, was able to almost completely alleviate the repression of the reporter activity by ZNF704 (Figure 2D, lower), suggesting that ZNF704-mediated transcription repression was associated with an HDAC activity.

In order to gain mechanistic insights into the transcription repression function of ZNF704, we employed affinity purification coupled with mass spectrometry to interrogate the ZNF704 interactome *in vivo*. In these experiments, FLAG-ZNF704 was stably expressed in MDA-MB-231 cells. Cellular extracts were prepared and subjected to affinity purification using an anti-FLAG affinity column, and the bound proteins were analyzed by mass spectrometry. The results showed that ZNF704 was co-purified with a series of proteins including SIN3A, SAP130, HDAC1, HDAC2, and RBBP4, all components of the SIN3A complex (Figure 2E, left). Additional proteins including PRKDC and DDB1 were also detected in the ZNF704-containing complex (Figure 2E, left). The presence of the SIN3A components in the ZNF704-associated protein complex was verified by western blotting of the column eluates (Figure 2E, right). The association between ZNF704 and the SIN3A complex was also detected in HEK293T cells by affinity purification-coupled mass spectrometry (Figure 2E, lower). The detailed results of the mass spectrometric analysis are provided in Supplemental Table S1. Together, these results indicate that ZNF704 is associated with the SIN3A transcription corepressor complex *in vivo*.

To verify the *in vivo* interaction between ZNF704 and the SIN3A corepressor complex, total proteins from MDA-MB-231 cells were extracted and co-immuoprecipitation was performed with antibodies detecting the endogenous proteins. Immunoprecipitation (IP) with antibodies against ZNF704 followed by immunoblotting (IB) with antibodies against the components of the SIN3A corepressor complex demonstrated that the constituents of the SIN3A corepressor complex were efficiently co-immunoprecipitated with ZNF704 (Figure 2F, left). Reciprocally, IP with antibodies against representative components of the SIN3A complex and IB with antibodies against ZNF704 also showed that ZNF704 was co-immunoprecipitated with the components of the SIN3A corepressor complex (Figure 2F, upper). In addition, the association between ZNF704 and the SIN3A corepressor complex was also detected in MCF-7 cells by co-immuoprecipitation assays (Figure 2F, right).

To further support the physical interaction of ZNF704 with the SIN3A corepressor complex and to understand the molecular basis underlying this interaction, glutathione S-transferase (GST) pull-down assays were performed with GST-fused ZNF704 (GST-ZNF704) and *in vitro* transcribed/translated individual components of the SIN3A corepressor complex. These experiments revealed that ZNF704 was capable of interacting with SIN3A and SAP130, but not with the other components of the SIN3A corepressor complex that we tested (Figure 2G), suggesting that the association of ZNF704 with the SIN3A corepressor complex is through its interactions with SIN3A and SAP130.

To further substantiate the physical interaction of ZNF704 with the SIN3A corepressor complex *in vivo*, nuclear proteins extracted in high salts from MDA-MB-231 cells were fractionated by size exclusion using fast protein liquid chromatography (FPLC) with Superose 6 column. We found that native nuclear ZNF704 from MDA-MB-231 extracts was eluted with an apparent molecular mass much greater than that of the monomeric protein (Figure 2H, left); ZNF704 immunoreactivity was detected in chromatographic fractions with an elution pattern that largely overlapped with that of the subunits of the SIN3A corepressor complex including SIN3A, SAP180, SAP130, HDAC1/2, and RBBP4/7 (Figure 2H, left). Importantly, analysis of the FLAG-ZNF704 affinity eluate from FPLC after Superose 6 gel filtration in MDA-MB-231 cells stably expressing FLAG-ZNF704 detected a multiprotein complex containing SIN3A, SAP180, and HDAC1/2 (Figure 2H, right). Collectively, these experiments support the observation that ZNF704 is physically associated with the SIN3A corepressor complex *in vivo*.

### Genome-wide Identification of the Transcriptional Targets for the ZNF704/SIN3A Complex

To explore the biological significance of the physical interaction between the transcription repressor ZNF704 and the SIN3A corepressor complex, we next analyzed the genome-wide transcriptional targets of the ZNF704/SIN3A complex. To this end, chromatin immunoprecipitation-based deep sequencing (ChIP-seq) was performed in MDA-MB-231 cells first using antibodies against ZNF704 or SIN3A. Following ChIP, ZNF704- and SIN3A-associated DNAs were amplified using non-biased conditions, labeled, and then sequenced via BGISEQ-500. With Model-based Analysis for ChIP-seq version 14 (MACS14) and a p value cutoff of 10^−3^, we identified 22,493 ZNF704-specific binding peaks and 16,576 SIN3A-specific binding summits (Figure 3A). The DNA sequences associated with these peaks were then cross-analyzed for overlapping gene promoters to represent the co-targets of ZNF704 and the SIN3A complex. These analyses identified a total of 1,354 promoters targeted by the ZNF704/SIN3A complex, which were then classified by gene ontology with DAVID (https://david.ncifcrf.gov/) into different KEGG pathways (Figure 3B). These KEGG pathways include hippo signaling, circadian rhythm, and MAPK signaling pathway that are well established to play important roles in tumorigenesis (Figure 3B). Significantly, analysis of the genomic signatures of ZNF704 and SIN3A revealed indeed similar binding motifs for these two proteins (Figure 3C), strongly supporting the physical interaction and functional connection between ZNF704 and SIN3A.

**Figure 3.**
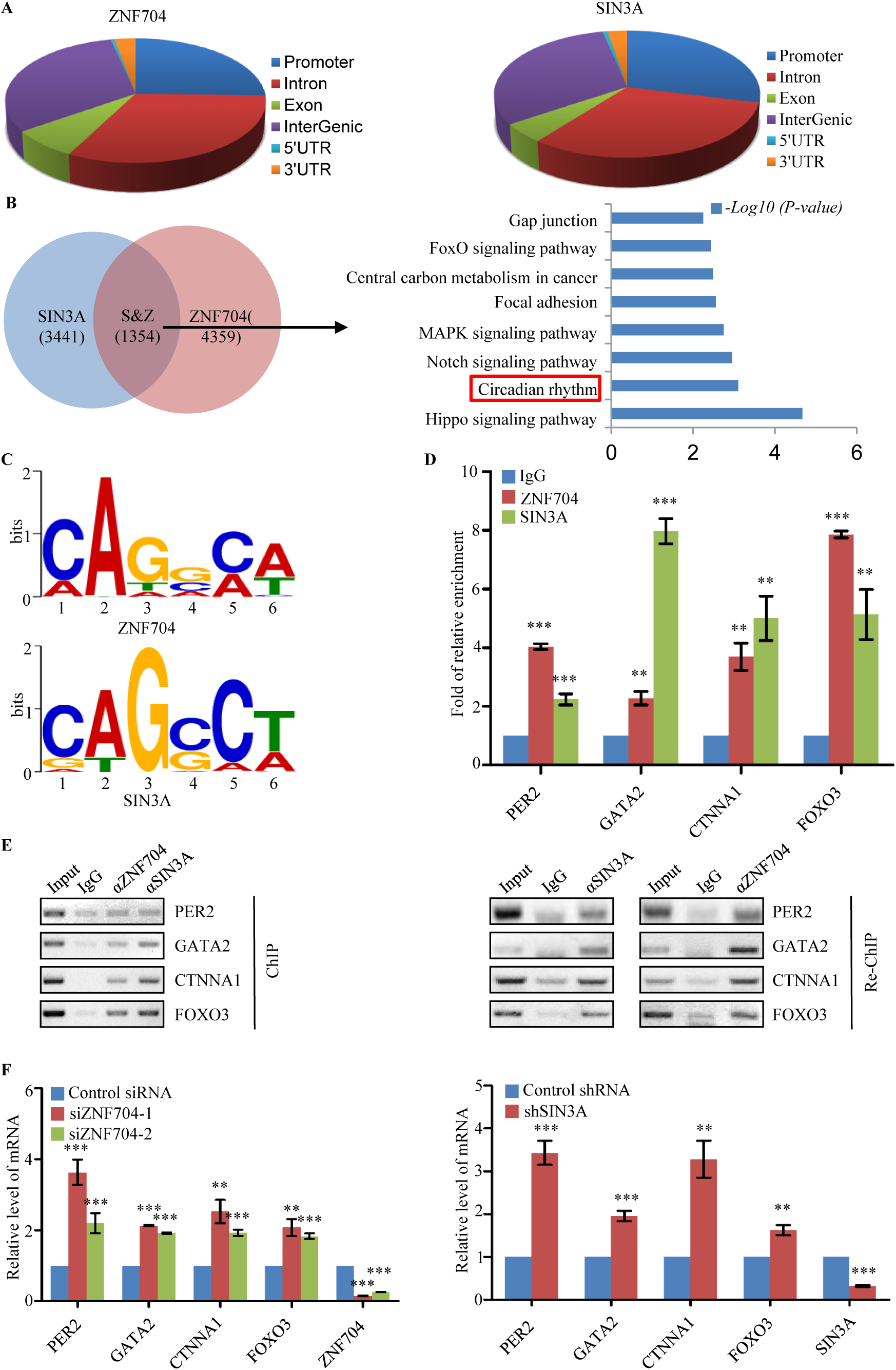
Genome-wide Identification of the Transcriptional Targets for the ZNF704/SIN3A Complex. (A) ChIP-seq analysis of the genomic distribution of the transcriptional targets of ZNF704 and SIN3A in MDA-MB-231 cells. (B) The overlapping genes targeted by ZNF704 and SIN3A in MDA-MB-231 cells (left). The results from KEGG analysis of co-targets are shown (right). (C) MEME analysis of the DNA-binding motifs of ZNF704 and SIIN3A. (D) qChIP verification of the ChIP-seq results on the promoter of the indicated genes with antibodies against ZNF704 and SIN3A in MDA-MB-231 cells. Results are presented as fold of change over control. Error bars represent mean ± SD for triplicate experiments. (E) ChIP/Re-ChIP experiments on the promoter of the indicated genes with antibodies against ZNF704 and SIN3A in MDA-MB-231 cells. (F) qPCR measurement of the expression of the indicated genes selected from ChIP-seq results in MDA-MB-231 cells under knockdown of ZNF704 or SIN3A. The knockdown efficiency was validated by qPCR. Error bars represent mean ± SD for triplicate experiments. Data information: In (D, F), data are presented as mean ± SEM. \**P* <0.05, \*\*\**P* < 0.001 (Student’s t-test).

ChIP-seq results were then validated by quantitative ChIP (qChIP) analysis in MDA-MB-231 cells using specific antibodies against ZNF704 or SIN3A on selected gene promoters including *PER2*, *GATA2*, *CTNNA1*, and *FOXO3.* The results showed a strong enrichment of ZNF704 and SIN3A on the promoters of these genes (Figure 3D). To verify that ZNF704 and SIN3A existed in the same protein complex on target gene promoters, we performed sequential ChIP or ChIP/Re-ChIP on representative target genes, *PER2*, *GATA2*, *CTNNA1*, and *FOXO3*. In these experiments, soluble chromatin was initially IP with antibodies against ZNF704, and the immunoprecipitates were subsequently re-IP with antibodies against SIN3A. The results of these experiments showed that the *PER2*, *GATA2*, *CTNNA1*, and *FOXO3* promoters that were IP with antibodies against ZNF704 could be re-IP with antibodies against SIN3A (Figure 3E). Similar results were obtained when the initial ChIP was carried out with antibodies against SIN3A (Figure 3E). Together, these results validated the targeting of *PER2*, *GATA2*, *CTNNA1*, and *FOXO3* by the ZNF704/SIN3A complex and support the coexistence of ZNF704 and SIN3A on the promoter of these genes.

To further consolidate the ChIP-seq results, ZNF704 was knocked down in MDA-MB-231 cells using two different sets of small interfering RNA and the expression of *PER2*, *GATA2*, *CTNNA1*, and *FOXO3* was analyzed by real time RT-PCR. ZNF704 knockdown resulted in a significant increase, albeit to a different extent, in the expression of all the tested genes (Figure 3F, left). The knockdown efficiency was verified by real-time RT-PCR (Figure 3F, left). Similarly, depletion of SIN3A was also associated with an increased expression of the tested genes (Figure 3F, right). Together, these results support our observations that ZNF704 and the SIN3A complex are physically associated and functionally connected to repress downstream target genes.

### ZNF704 Transcriptionally Represses *PER2* and Functionally Disrupts Circadian Rhythm in Breast Cancer Cells

The identification of *PER2* as a target of the ZNF704/SIN3A complex suggests that the ZNF704/SIN3A complex might influence circadian clock in breast cancer cells. To test this, the effect of the ZNF704/SIN3A complex on the expression of PER2 protein was examined first in MDA-MB-231 cells transfected with lentivirally delivered vector or FLAG-ZNF704, and/or treated with lentivirally delivered scrambled short hairpin RNA (SCR shRNA) or shRNA against ZNF704 or SIN3A. Western blotting showed that ZNF704 overexpression led to a decrease in the level of PER2, which could be rescued by depletion of SIN3A (Figure 4A, left), whereas in ZNF704-depleted cells, the level of PER2 increased (Figure 4A, right).

**Figure 4.**
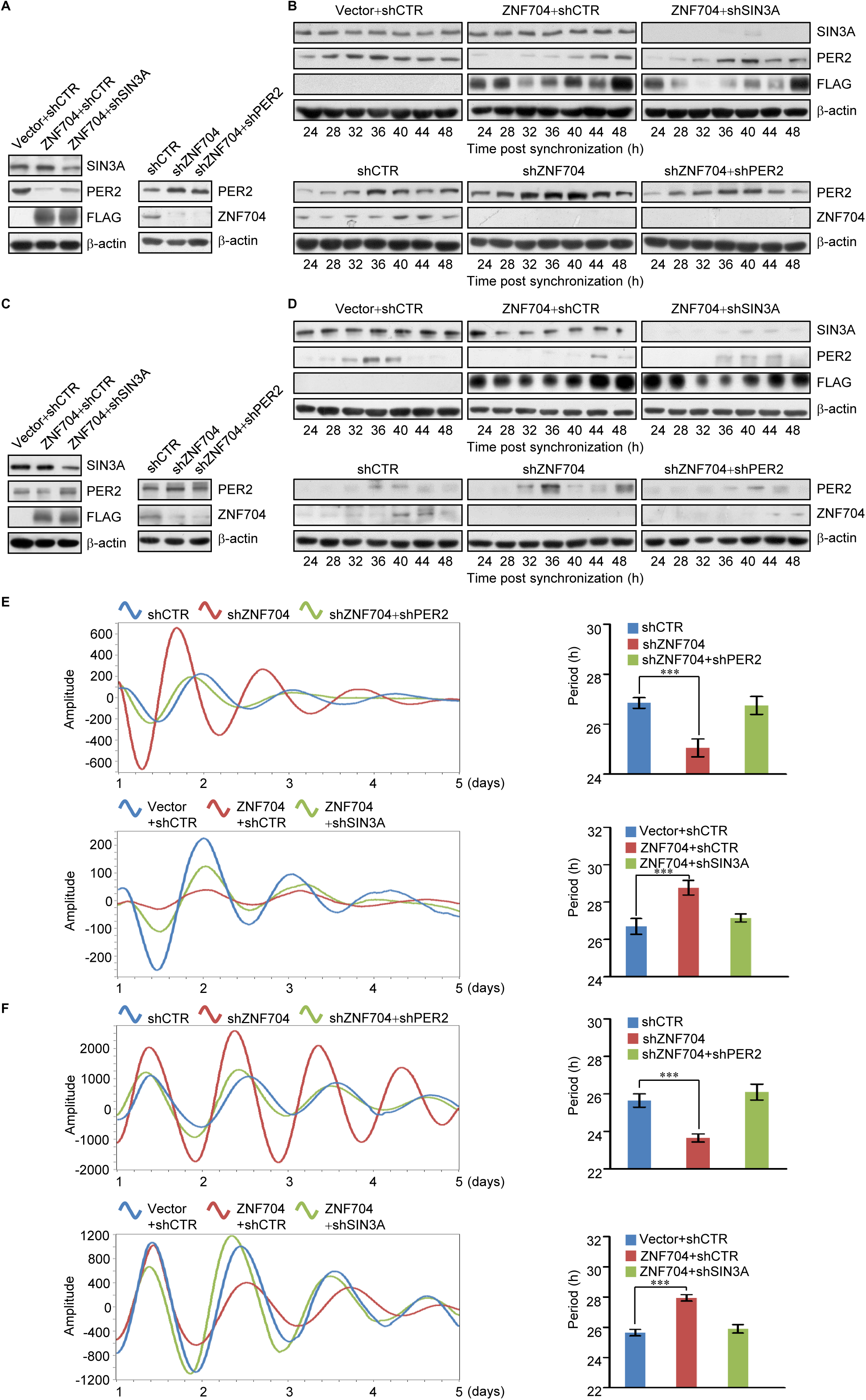
ZNF704 Transcriptionally Represses *PER2* and Functionally Disrupts Circadian Rhythm in Breast Cancer Cells. (A) MDA-MB-231 cells were infected with lentiviruses carrying the indicated expression constructs and/or specific shRNAs for the measurement of SIN3A, PER2, FLAG and ZNF704 by western blotting. (B) MDA-MB-231 cells infected with lentiviruses carrying the indicated expression constructs and/or specific shRNAs were collected at 4 h-interval from 24 to 48 h for the measurement of SIN3A, PER2, and ZNF704 by western blotting. (C) U2OS cells were infected with lentiviruses carrying the indicated expression constructs and/or specific shRNAs for the measurement of SIN3A, PER2, FLAG and ZNF704 by western blotting. (D) U2OS cells infected with lentiviruses carrying the indicated expression constructs and/or specific shRNAs were collected at 4 h-interval from 24 to 48 h for the measurement of SIN3A, PER2, FLAG and ZNF704 by western blotting. (E) MDA-MB-231*-Per2-dLuc* cells were infected lentiviruses carrying the indicated expression constructs and/or specific shRNAs for luciferase reporter assays (left). Histogram shows the quantitative period changes (right). Error bars represent mean ± SD for triplicate experiments. (F) U2OS*-Per2-dLuc* cells were infected lentiviruses carrying the indicated expression constructs and/or specific shRNAs for luciferase reporter assays (left). Histogram shows the quantitative period changes (right). Error bars represent mean ± SD for triplicate experiments. Data information: In (E-F), data are presented as mean ± SEM. \*\*\**P* < 0.001 (Student’s t-test).

To further investigate the influence of the ZNF704/SIN3A complex on the oscillation of PER2 protein expression, MDA-MB-231 cells that were transfected with vector or FLAG-ZNF704, and/or SCR shRNA or shRNA against ZNF704 or SIN3A were synchronized by serum starvation for 24 h followed by treatment with dexamethasone for 2 h. The cells were then switched to serum-free media and collected at a 4-h interval. Western blotting analysis revealed that overexpression of ZNF704 inhibited the baseline of PER2 level and altered the oscillation of PER2 expression, effects that could be at least partially counteracted by depletion of SIN3A (Figure 4B, upper). Conversely, knockdown of ZNF704 resulted in an increase in the baseline of PER2 level and also altered oscillation of PER2 expression, which could be rescued by simultaneous knockdown of PER2 (Figure 4B, lower). Similar effects on PER2 expression (Figure 4C) and oscillation (Figure 4D) were also obtained in U2OS cells, which were used as the model system for circadian rhythm study(Baggs et al, 2009; Hirota et al, 2008; Maier et al, 2009).

To gain further insight into the effect of the ZNF704/SIN3A complex on circadian rhythm, MDA-MB-231 cells that stably express *Per2* promoter-driven luciferase were generated. These cells were transfected with lentivirally delivered vector or FLAG-ZNF704, and/or treated with lentivirally delivered SCR shRNA or shRNA against ZNF704, PER2, or SIN3A. Monitoring the cells with real-time Lumi-Cycle luminometry showed that knockdown of ZNF704 led to a period-shortening and amplitude-increasing phenotype, effects that were at least partially attenuated by co-knockdown of PER2 (Figure 4E, upper). Conversely, we observed a period-lengthening and amplitude-damping phenotype when ZNF704 was overexpressed, and this effect could be rescued, at least partially, by simultaneous depletion of SIN3A (Figure 4E, lower). Similar results were also obtained in U2OS cells (Figure 4F). Together, these observations indicate that ZNF704 overexpression disrupts the circadian rhythm, and that ZNF704 does so, through cooperating with the SIN3A complex to repress PER2 expression.

### The ZNF704/SIN3A Complex Promotes the Proliferation and Invasion of Breast Cancer Cells *in Vitro*

Dysfunction of circadian rhythm is closely associated with the development and progression of various malignancies including breast cancer(Hoffman et al, 2010a; Hoffman et al, 2010b; Iurisci et al, 2010; Kochan & Kovalchuk, 2015). Likewise, *PER2* is also implicated in tumorigenesis and has been proposed as a tumor suppressor(Fu et al, 2002; Hwang-Verslues et al, 2013; Sun et al, 2010; Wang et al, 2016). In light of the observations that ZNF704 represses the expression of PER2 and ZNF704 overexpression leads to the disruption of circadian rhythm, it is reasonable to postulate that ZNF704 could affect the development and progression of breast cancer. To test this, gain-of-function and loss-of-function experiments of ZNF704 were performed in MCF-7 cells and the effect of ZNF704 on the proliferation of these cells was examined using CCK-8 (Cell Counting Kit-8) assays. The results showed that ZNF704 overexpression promoted breast cancer cell proliferation, an effect that could be abrogated, at least partially, by knockdown of SIN3A (Figure 5A, left). Consistently, depletion of ZNF704 had a significant inhibitory effect on breast cancer cell proliferation, a phenotype that could be rescued by co-knockdown of PER2 (Figure 5A, right). Moreover, colony formation assays in MCF-7 cells showed that overexpression of ZNF704 resulted in an increased in colony number, which was abrogated upon depletion of SIN3A (Figure 5B, upper), whereas knockdown of ZNF704 was associated with a decreased colony number, a phenotype that could be, at least partially, rescued by co-knockdown of PER2 (Figure 5B, lower). Together, these experiments support a role for ZNF704 in promoting breast cancer cell proliferation, indicating that ZNF704 does so, through association with the SIN3A complex and downregulation of target genes including *PER2*.

**Figure 5.**
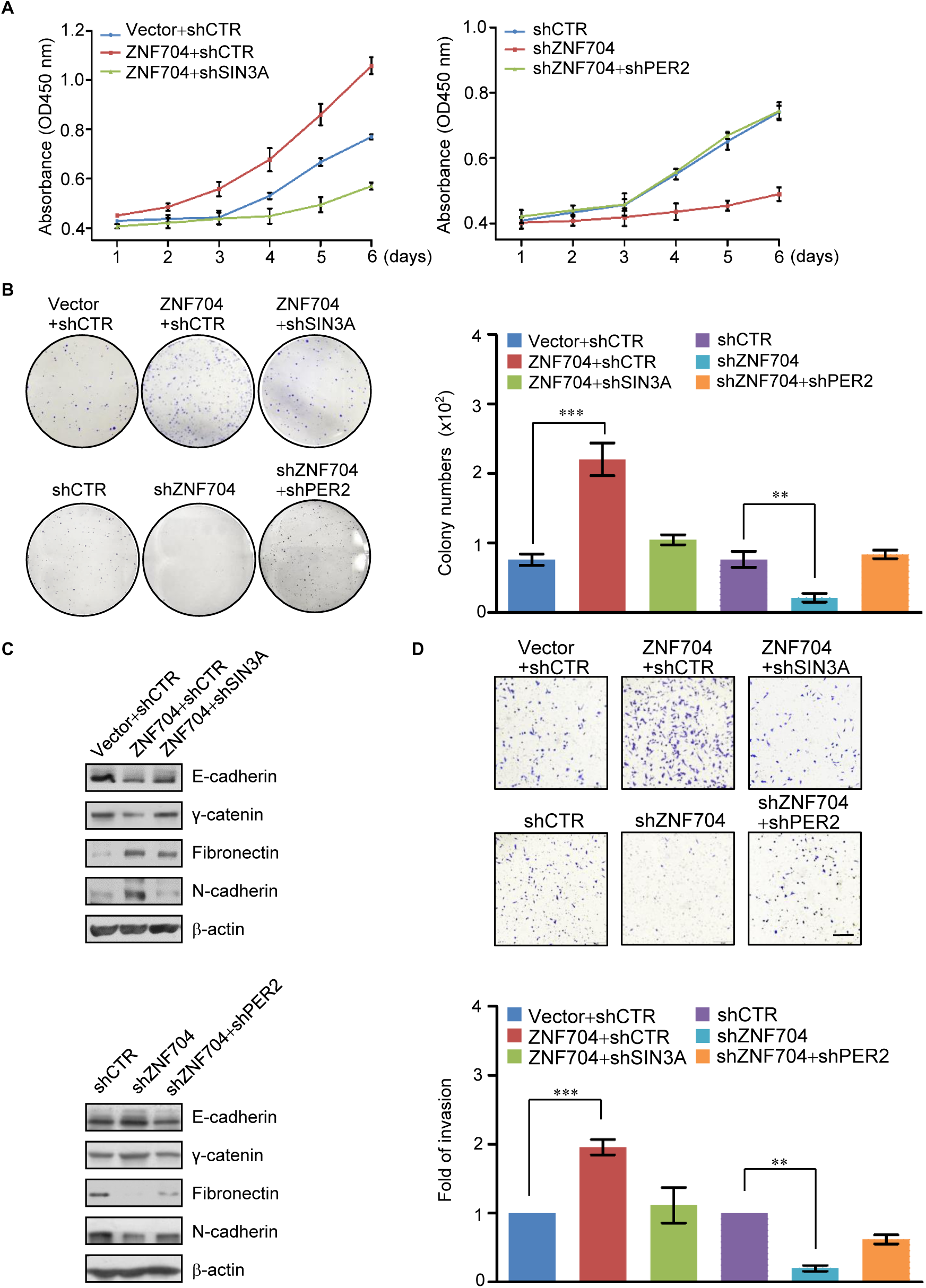
The ZNF704/SIN3A Complex Promotes the Proliferation and Invasion of Breast Cancer Cells *in Vitro*. (A) CCK-8 assays for the proliferation of MCF-7 cells infected with lentiviruses carrying the indicated expression constructs and/or specific shRNAs. Error bars represent the mean ± SD for three independent experiments. (B) MCF-7 cells infected with lentiviruses carrying the indicated expression constructs and/or specific shRNAs were cultured for 14 days before staining with crystal violet and counting for colony numbers. Error bars represent the mean ± SD for three independent experiments. (C) MDA-MB-231 cells were infected with lentiviruses carrying the indicated expression constructs and/or specific shRNAs for the measurement of the expression of the indicated epithelial/mesenchymal markers by western blotting. (D) MDA-MB-231 cells were infected with lentiviruses carrying the indicated expression constructs and/or specific shRNA for transwell invasion assays. The invaded cells were stained and counted. The images represent one microscope field in each group. Error bars represent mean ± SD for triplicate experiments. Bar: 50 μm. Data information: In (B, D), data are presented as mean ± SEM. \*\**P* < 0.01, \*\*\**P* < 0.001 (Student’s t-test).

To investigate the role of ZNF704 in breast cancer progression, the expression of epithelial/mesenchymal markers was first analyzed by western blotting in MDA-MB-231 cells, as epithelial-mesenchymal transition (EMT) is believed to be the initial step of tumor metastasis(Wang & Shang, 2013). We found that overexpression of ZNF704 resulted in a reduction of epithelial markers including E-cadherin and γ-catenin and an induction of mesenchymal markers including fibronectin and N-cadherin, which were at least partially attenuated by co-knockdown of SIN3A (Figure 5C, upper). Conversely, depletion of ZNF704 was associated with an induction of the epithelial markers and reduction of the mesenchymal markers (Figure 5C, lower). However, simultaneous depletion of PER2 counteracted the effect of ZNF704 depletion on the expression patterns of the epithelial/mesenchymal markers (Figure 5C, lower). These results support a role for ZNF704 in promoting EMT.

We then investigated the role of ZNF704 in the cellular behavior of breast cancer cells *in vitro* using transwell invasion assays. We found that ZNF704 overexpression was associated with an increase in the invasive potential of MDA-MB-231 cells, whereas ZNF704 knockdown was accompanied by a decrease in the invasive potential of MDA-MB-231 cells (Figure 5D). Moreover, in agreement with the functional link between ZNF704 and SIN3A, the increase in invasive potential associated with ZNF704 overexpression could be offset, at least partially, by knockdown of SIN3A and the inhibitory effect of ZNF704 knockdown on the invasive potential of MDA-MB-231 cells was at least partially rescued by PER2 depletion (Figure 5D). Taken together, these results indicate a role for ZNF704 in regulating the invasive potential of breast cancer cells and support the functional link between the ZNF704/SIN3A complex and PER2.

### The ZNF704/SIN3A Complex Promotes the Growth and Metastasis of Breast Cancer *in Vivo*

To investigate the role of ZNF704 in breast cancer metastasis *in vivo*, MDA-MB-231 cells that had been engineered to stably express firefly luciferase (MDA-MB-231-Luc-D3H2LN, Xenogen Corporation) were infected with lentiviruses carrying vector or FLAG-ZNF704, or/and carrying shCTR or shRNAs against ZNF704, SIN3A, or PER2. These cells were then implanted onto the left abdominal mammary fat pad of 6-week-old female SCID mice (n = 6). The growth/dissemination of tumors was monitored weekly by bioluminescence imaging with IVIS imaging system. Tumor metastasis was measured by quantitative bioluminescence imaging after 6 weeks. A metastatic event was defined as any detectable luciferase signal above background and away from the primary tumor site. The results showed that ZNF704 overexpression promoted the growth of the primary tumor and accelerated the lung metastasis of the MDA-MB-231-Luc-D3H2LN tumors (Figure 6A). However, depletion of SIN3A neutralized the ZNF704 overexpression-associated promoting effects of the growth of primary tumors and lung metastases (Figure 6A). Consistently, ZNF704 depletion resulted in inhibition of the growth of the primary tumor and suppression of the lung metastasis of the MDA-MB-231-Luc-D3H2LN tumors, effects that could be offset by co-knockdown of PER2 (Figure 6A).

**Figure 6.**
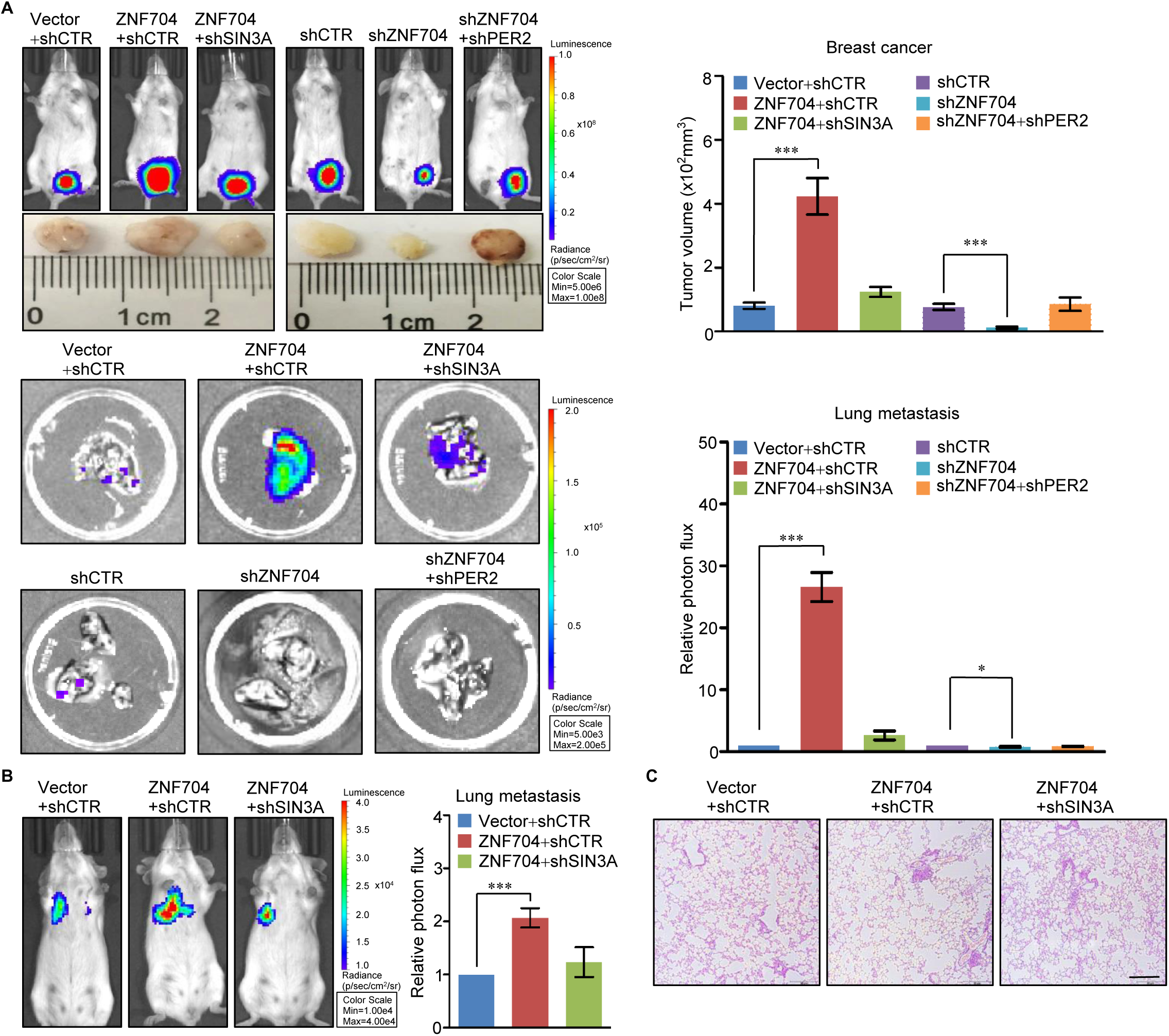
The ZNF704/SIN3A Complex Promotes the Growth and Metastasis of Breast Cancer *in Vivo*. (A) MDA-MB-231-Luc-D3H2LN cells infected with lentiviruses carrying the indicated expression constructs and/or specific shRNA were inoculated orthotopically onto the abdominal mammary fat pad of 6-week-old female SCID mice (n = 6). Primary tumor size was measured using bioluminescent imaging after 6 weeks of initial implantation. Representative primary tumors and bioluminescent images are shown. Error bars represent mean ± SD for three independent measurements. (B) MDA-MB-231 Luc-D3H2LN cells infected with lentiviruses carrying the indicated expression constructs and/or specific shRNAs were injected intravenously into 6-week-old female SCID mice (n = 6). Metastases were quantified using bioluminescence imaging after 4 weeks of initial implantation. Representative bioluminescent images are shown. Error bars represent mean ± SD for three independent measurements. (C) Representative lung metastasis specimens were sectioned and stained with H&E. Bar: 50 μm. Data information: In (A, B), data are presented as mean ± SEM. \**P* < 0.05, \*\**P* < 0.01, \*\*\**P* < 0.001 (Student’s t-test).

Next, MDA-MB-231 Luc-D3H2LN cells infected with lentiviruses carrying vector or FLAG-ZNF704 or/and carrying shCTR or shRNA against SIN3A were injected intravenously into SCID mice (n = 6), and seeding lung metastasis was measured by quantitative bioluminescence imaging after 4 weeks of injection. The results showed that overexpression of ZNF704 drastically promoted lung metastasis of the MDA-MB-231-Luc-D3H2LN tumors, and this was attenuated at least partially by simultaneous knockdown of SIN3A (Figure 6B). The lung metastasis was verified by histological staining (Figure 6C). Collectively, these experiments indicate that ZNF704 promotes the growth and metastasis of breast cancer, and that it does so, through its interaction with the SIN3A complex and repression of target genes including *PER2*.

### High Level of ZNF704 Is Correlated with Worse Clinical Behaviors and Poor Prognosis of Breast Cancer Patients

To extend our observations to clinicopathologically relevant contexts, we collected 25 breast carcinoma samples paired with adjacent normal mammary tissues from breast cancer patients and analyzed by qPCR for the expression of ZNF704 and PER2. We found that the mRNA level of ZNF704 is upregulated, whereas the mRNA level of PER2 is downregulated in these breast carcinoma samples (Figure 7A). In line with our working model that ZNF704 and its associated SIN3A corepressor complex transcriptionally repress PER2, when the relative mRNA levels of PER2 were plotted against that of ZNF704 in the 25 breast carcinoma samples, a significant negative correlation was found (Figure 7B). In addition, querying published clinical datasets (GSE27562 and GSE3744) showed a clear negative correlation of the mRNA levels between ZNF704 and PER2 (Figure 7C). Moreover, interrogation of Lu’s breast cancer dataset (Figure 7D) in Oncomine (https://www.oncomine.org/) as well as the public dataset (GSE61304) (Figure 7E) showed that the level of ZNF704 expression is negatively correlated with the histological grades of breast cancer. Furthermore, analysis of the public dataset (GSE36774) found that high ZNF704 and low PER2 in breast carcinomas strongly correlated with lymph node positivity of the patients (Figure 7F), and, remarkably, analysis of the public dataset (GSE65194) revealed that the level of ZNF704 expression is higher in HER2-enriched breast carcinomas than in luminal A and B subtypes, whereas the level of PER2 expression exhibited a reverse trend (Figure 7G).

**Figure 7.**
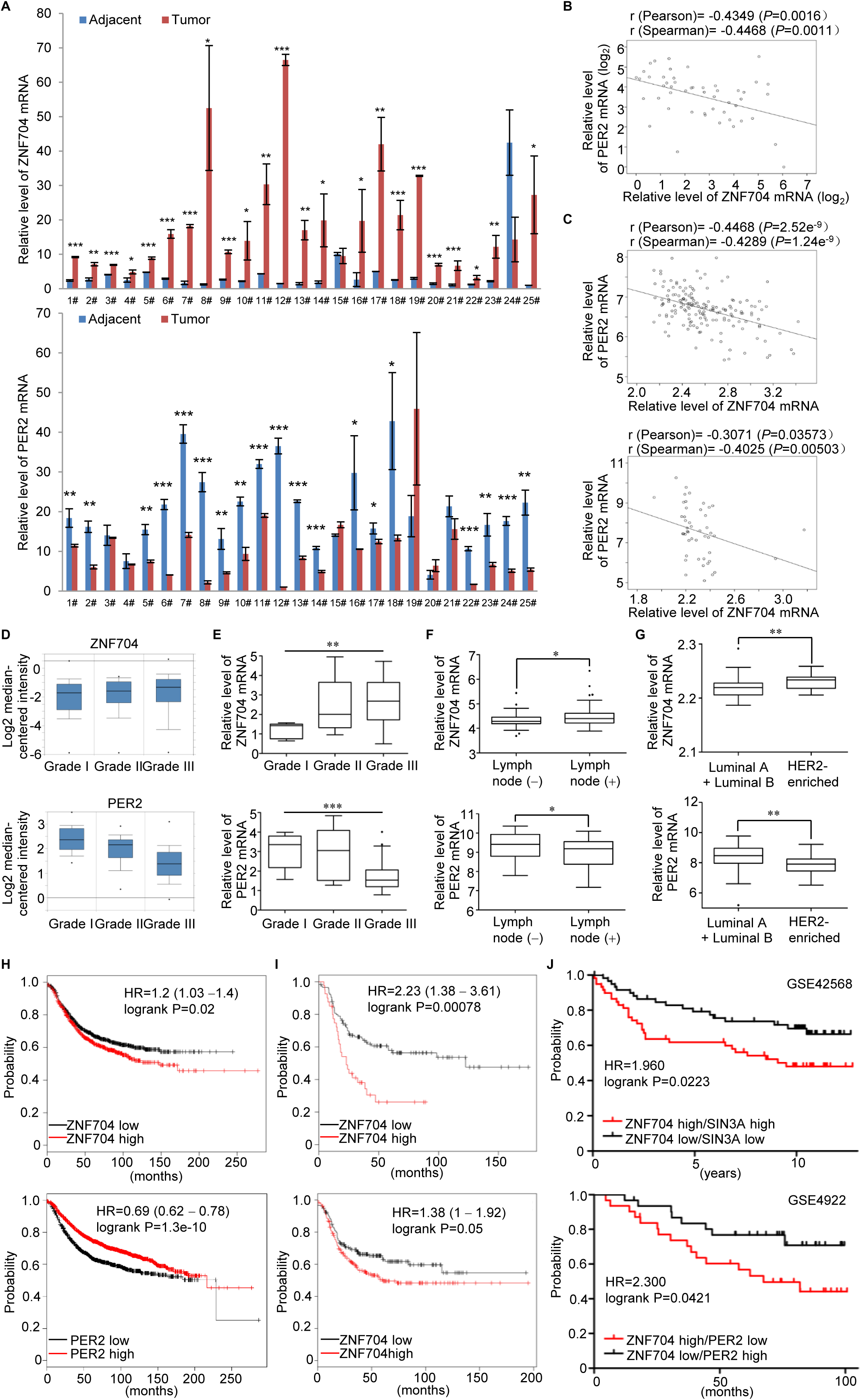
High Level of ZNF704 Is Correlated with Worse Clinical Behaviors and Poor Prognosis of Breast Cancer Patients. (A) Analysis of ZNF704 and PER2 expression by real-time RT-PCR in 25 breast carcinoma samples paired with adjacent normal mammary tissues. Each bar represents the mean ± SD for triplicate experiments. (B) The relative level of ZNF704 was plotted against that of PER2. The correlation coefficients were calculated by SPSS19.0. (C) The relative level of ZNF704 was plotted against that of PER2 based on public datasets GSE27562 (upper) and GSE3744 (lower). (D) The correlation between ZNF704 or PER2 expression and histological grade in Lu’s breast cancer dataset from Oncomine (https://www.oncomine.org/). (E) The correlation between ZNF704 or PER2 expression and histological grade in public dataset (GSE61304). (F) Analysis of public dataset (GSE36774) for the correlation between the level of ZNF704 or PER2 and the lymph node metastasis of breast cancer patients. (G) Analysis of public dataset (GSE65194) for the correlation between the level of ZNF704 or PER2 and the molecular subtypes of breast cancer patients. (H) Kaplan-Meier survival analysis for the relationship between survival time and ZNF704 (upper) or PER2 (lower) signature in breast cancer using the online tool (http://kmplot.com/analysis/). (I) Kaplan–Meier survival analysis of the published datasets for the relationship between survival time and ZNF704 signature in HER2-enriched (upper) or basal-like (lower) breast cancer using the online tool (http://kmplot.com/analysis/). (J) Kaplan–Meier survival analysis of the published datasets (GSE42568 and GSE4922) for the relationship between survival time and ZNF704/PER2 or ZNF704/SIN3A signature in breast cancer. Data information: In (A, F, G), data are presented as mean ± SEM. \**P* < 0.05, \*\**P* < 0.01, \*\*\**P* < 0.001 (Student’s t-test); In (E), data are presented as mean ± SEM. **P < 0.01, *** P < 0.001 (one-way ANOVA).

Finally, Kaplan-Meier survival analysis (http://kmplot.com/analysis/) of public datasets found that either higher ZNF704 expression (hazard ratio, HR=1.2, *P*=0.02) or lower PER2 expression (HR=0.69, *P*=1.3e-10) was associated with a poorer relapse-free survival of breast cancer patients, when the influence of systemic treatment, endocrine therapy, and chemotherapy were excluded (Figure 7H). This is particularly true for HER2-enriched and basal-like subtypes of breast cancer patients (Figure 7I). Further analysis of the public datasets (GSE42568 and GSE4922) by stratifying patient groups based on inverse expression patterns of ZNF704 and PER2 or the co-expression of ZNF704 and SIN3A significantly improved the predictive capability of ZNF704 (Figure 7J). Collectively, these analyses support our observations that ZNF704 is a transcription repressor and a potent driver of breast cancer development and progression.

## Discussion

Gene amplification is an important mechanism for protein overexpression and oncogene hyperactivation in tumorous cells(Myllykangas et al, 2007). Although amplification/copy number gains at 8q21 is a frequent event in various malignancies including breast cancer, a genetic alteration often associated with poor prognosis of the patients(Balleine et al, 2000; Choschzick et al, 2010), the genetic factor(s) that potentially contribute to the oncogenic potential of the 8q21 amplicon remains to be determined. In this study, we found *ZNF704*, a gene that is mapped to 8q21 and encodes for a zinc finger transcription factor, is recurrently amplified in breast cancer and other types of cancer. We showed that ZNF704 acts as a transcription repressor and the transcriptional repression activity of ZNF704 is associated with a histone deacetylase activity. Indeed, immunopurification-coupled mass spectrometry demonstrated that ZNF704 is associated with the SIN3A complex, a multi-protein assembly containing HDAC1/HDAC2. The SIN3A complex has been extensively studied as a corepressor complex that is recruited by a number of transcription factors and functions in a panel of biological activities including embryonic development(Streubel et al, 2017), stem cell differentiation(Mcdonel et al, 2012), and tumor progression(Shan et al, 2016). Our finding that this complex is functionally involved in gene(s) residing in the 8q21 amplicon is consistent with the role of the SIN3A complex in tumorigenesis. Interestingly, genome-wide interrogation of the transcriptional targets by ChIP-seq identified that the ZNF704/SIN3A complex represses a cohort of genes including *PER2* that is an essential component that constitutes the molecular system controlling the circadian clock.

Dysfunction of circadian clock has been linked to the development and progression of tumors(Altman et al, 2014; Yeh et al, 2014; Yu & Weaver, 2011), yet the regulation and deregulation of core clock genes in tumorigenesis is less understood. Among the clock genes, *PER2* dysregulation or deletion is also frequently observed in malignancies from a broad spectrum of tissue origins, and these aberrations are associated with a more aggressive phenotype and accordingly poorer survival of the cancer patients(Liu et al, 2014; Xiong et al, 2018; Zhao et al, 2014). Of note, *PER2* promoter hypermethylation is detected in endometrial cancer(Shih et al, 2010) and glioma(Fan et al, 2014), suggesting that transcriptional regulation of PER2 is pathologically relevant to tumor development and progression. Our study showed that PER2 is transrepressed by the ZNF704/SIN3A complex. We demonstrated that overexpression of ZNF704 prolongs the period and dampens the amplitude of circadian clock in breast cancer cells. Moreover, ZNF704 overexpression promotes the proliferation and invasion of breast cancer cells *in vitro* and facilitates the growth and metastasis of breast cancer *in vivo*. It is conceivable that in breast cancer cells, accompanying with 8q21 amplification, *ZNF704* is amplified and ZNF704 is overexpressed, which, in turn, down-regulates PER2, leading to the disruption of circadian rhythm, eventually contributing to breast carcinogenesis.

The transcription regulation of PER2 by ZNF704 is interesting. After all, the consensus is that the molecular clock is driven by a systemic feedback loop, in which CLOCK-BMAL1 induces PER and CRY proteins, and these proteins in turn form inhibitory complexes with CLOCK-BMAL1 to repress their own expression(Hida et al, 2000; Kume et al, 1999). Nevertheless, it is reported that CLOCK-BMAL1 also induces REV-ERBα and REV-ERBβ, which transcriptionally repress BMAL1 at retinoic acid receptor-related orphan receptor elements (ROREs), thereby constituting a second interlocking feedback loop(Preitner et al, 2002). Moreover, it was found that *PER* and *CRY* genes could still tick even with the depletion of CLOCK or the repression of BMAL1(Debruyne et al, 2006; Kornmann et al, 2007). These observations suggest a more complex molecular clock system and indicate that additional regulators exist that are critically involved in the regulation of circadian clock. Whether or not ZNF704 represents one of the additional regulators under physiological conditions remains to be investigated, and the functional relationship between ZNF704 and the CLOCK-BMAL1 heterodimer remains to be delineated. Perhaps more relevant to our study, whether and how ZNF704 exerts its oncogenic role in related to other oncogenic factors, especially the genes located at the 8q21 amplicon, await for future investigations. In these regards, it is important to note that additional genes implicated in several key cellular processes including cell proliferation, migration, and molecular catabolism were also identified to be the transcriptional targets of the ZNF704/SIN3A complex. Although the multitude of the cellular function of the ZNF704 is probably beyond the scope of our current investigation, we nevertheless don’t exclude the possibility of other transcriptional targets of ZNF704 in assisting or in contributing to breast carcinogenesis or the development and progression of cancers from other tissue origins.

In support of the role of ZNF704 in promoting breast carcinogenesis, we found that ZNF704 is highly expressed in breast cancer samples, and in agreement with our working model that ZNF704 enlists the SIN3A complex to repress *PER2* in its oncogenic activity, we found that the level of ZNF704 is negatively correlated with that of PER2, and we showed that high level of ZNF704 correlates with advanced histological grades and lymph node positivity of breast carcinomas and poor prognosis of breast cancer patients, especially those with HER2^+^ and basal-like subtypes. More studies are needed to gain mechanistic insights into the association of ZNF704 with particular subtypes of breast cancer and to evaluate whether these observations might yield potential prognostic values for breast cancer.

In summary, we report in the current study that ZNF704 is physically associated with the SIN3A complex and functionally coordinates histone deacetylation to repress downstream target genes including *PER2* to disrupt circadian rhythm to promote breast carcinogenesis. These observations indicate a critical role for ZNF704 in breast carcinogenesis, supporting the pursuit of ZNF704 as a therapy target for breast cancer intervention.

## Methods and materials

### Antibodies and Reagents

The cDNA for wild-type of ZNF704 was amplified by PCR and ligated into Xho I/Xba I sites of pCMV-IRES vector that contains three copies of FLAG. The GST-ZNF704 expression plasmid was constructed into a pGEX-4T-3 vector. All clones were confirmed by DNA sequencing. pLV7-Bsd-P(Per2)-KB-dLuc vector was a gift from Dr. Andrew C. Liu (Department of Biological Sciences, the University of Memphis, USA). Antibodies used were: αZNF704 from Beijing Protein Innovation Company; αSAP180 (24499-1-AP), αE-cadherin (20874-1-AP), αN-cadherin (22018-1-AP), and αfibronectin (15613-1-AP) from Proteintech; αSIN3A (GTX129156) from GeneTex; SAP130 (A302-490A) from Bethyl Laboratories; α-catenin (610253) from BD biosciences; αPER2 (ab179813) from Abcam; αHDAC1 (H3284), αHDAC2 (H3159), αRBBP4/7 (R3779), and αFLAG (F3165) from Sigma-Aldrich; αβ-actin (AC004) from Abclonal; The specificity of the bands in western blots was controlled according to the commercial antibodies’ instructions. Anti-FLAG M2 affinity gel (A2220) and 3 x FLAG peptide (F3290) were from Sigma-Aldrich, protein A or G beads were from Invitrogen (10004D), and protease inhibitor cocktail was from Roche Applied Science.

### Cell Culture and Transfection

Cell lines used were obtained from the American Type Culture Collection (ATCC). 293T, HeLa, U2OS and MCF-7 cells were maintained in DMEM supplemented with 10% FBS in a humidified incubator equilibrated with 5% CO_2_ at 37°C. MDA-MB-231 cells were cultured in L-15 medium supplemented with 10% FBS without CO_2_. Transfections were carried out using Polyethyienimine (Polysciences) or Lipofectamine RNAiMAX Reagent (Invitrogen) according to the manufacturer’s instructions. Each experiment was performed in triplicate and repeated at least three times. For RNAi experiment, at least three independent siRNA/shRNA sequences were tested for each gene, and the one with the best efficiency was used. The sequences of siRNA were: control siRNA, 5′-UUCUCCGAACGUGUCACGU-3′; ZNF704 siRNA-1, 5′-CAAUGGUACUAACCAGCUUGU-3′; ZNF704 siRNA-2, 5′-CCCUUUGGUUCGAAGUCCU-3′; Control siRNA and siRNAs for ZNF704 were synthesized by Sigma-Aldrich. The siRNA oligonucleotides were transfected into cells using RNAiMAX with a final concentration of 20 nM.

### Lentiviral Production and Infection

The generation of IRES-ZNF704, pLKO.1-shZNF704, pLKO.1-shPER2, or pLKO.1-shSIN3A lentiviruses was conducted according to a protocol described by Addgene. Briefly, human expression plasmid of IRES-ZNF704 was generated by subcloning ZNF704 cDNA into pCMV-IRES vector, and pLKO.1-shZNF704, pLKO.1-shPER2, and pLKO.1-shESIN3A were generated by subcloning shRNA (TRCN0000162553, shZNF704; TRCN0000330732, shPER2, TRCN0000162553, shSIN3A) into pLKO.1 vector. The lentiviral plasmid vector, pCMV-IRES, IRES-ZNF704, pLKO.1, pLKO.1-shZNF704, pLKO.1-PER2, or pLKO.1-shSIN3A, together with psPAX2 and pMD2.G, were co-transfected into the packaging cell line HEK293T. Viral supernatants were collected 48 h later, clarified by filtration, and concentrated by ultracentrifugation. The generation of pLV7-Bsd-P(Per2)-KB-dLuc lentiviruses was conducted according to the procedure described previously(Ramanathan et al, 2012). The concentrated viruses were used to infect 5 x 10^5^ cells (20-30% confluence) in a 60-mm dish with 5 μg/ml polybrene. Infected cells were selected by 2 μg/ml puromycin (Sigma) and/or hygromycin (Invitrogen) or blasticidin (Abcam). For re-silencing PER2 or SIN3A experiments, the level of PER2 or SIN3A expression was controlled by creating stable clones of cells that were expressing different levels of PER2 or SIN3A, and the clones with PER2 or SIN3A levels close to original PER2 or SIN3A levels were chosen for phenotype experiments.

### Fluorescence Confocal Microscopy

MCF-7 cells growing on 6-well chamber slides were washed with PBS, fixed in 4% (w/v) paraformaldehyde, permeabilized with 0.1% (v/v) Triton X-100 in PBS, blocked with 0.8% BSA, and incubated with appropriate primary antibodies followed by staining with ALEXA FLUOR-488/594 donkey or rabbit anti-mouse secondary antibodies (Invitrogen). The cells were washed 4 times and a final concentration of 0.1 μg/ml DAPI (Sigma) was included in the final washing to stain nuclei. Images were visualized with an Olympus inverted microscope equipped with a charge-couple camera.

### Luciferase Reporter Assay

Luciferase activity was measured using the Dual-Luciferase kit (Promega) according to the manufacturer’s protocol. Briefly, HeLa cells in 24-well plates were transfected with luciferase reporter, renilla, and appropriate expression constructs. The amount of DNA in each transfection was kept constant by addition of empty vector. Thirty-six hours after transfection, firefly and renilla luciferases were assayed according to the manufacturer’s protocol (Promega), and the activity of firefly luciferase was normalized to that of renilla luciferase. Each experiment was performed in triplicate and repeated at last three times.

### Silver Staining and Mass Spectrometry

MDA-MB-231 or 293T cells expressing FLAG-ZNF704 were washed twice with cold PBS, scraped, and collected by centrifugation at 800×g for 5 min. Cellular extracts were prepared by incubating the cells in lysis buffer containing protease inhibitor cocktail (Roche). Anti-FLAG immunoaffinity columns were prepared using anti-FLAG M2 affinity gel (Sigma) following the manufacturer’s suggestions. Cell lysates were obtained from about 5 x 10^8^ cells and applied to an equilibrated FLAG column of 1-ml bed volume to allow for adsorption of the protein complex to the column resin. After binding, the column was washed with cold PBS plus 0.1% Nonidet P-40 prior to application of 3 x FLAG peptides to elute FLAG protein complex as described by the vendor. Fractions of the bed volume were collected and resolved on NuPAGE 4-12% Bis-Tris gel (Invitrogen), silver-stained using Pierce silver stain kit, and subjected to LC-MS/MS (Agilent 6340) sequencing.

### Immunoprecipitation and Western Blotting

Cellular extracts from MDA-MB-231 or MCF-7 were prepared by incubating the cells in lysis buffer (50 mM Tris-HCl, pH8.0, 150 mM NaCl, 0.5% NP-40) for 30 min at 4°C. This was followed by centrifugation at 13,000 rpm for 15 min at 4°C. For immunoprecipitation, 500 μg of protein was incubated with specific antibodies (2-3 μg) for 12 h at 4°C with a constant rotation, and 30 μl of 50% protein A or G magnetic beads was then added and the incubation was continued for an additional 2 h. Beads were then washed three times using the lysis buffer. The precipitated proteins were eluted from the beads by resuspending the beads in 2 x SDS-PAGE loading buffer and boiling for 10 min. The resultant materials from immunoprecipitation or cell lysates were resolved using 10% SDS-PAGE gels and transferred onto acetate cellulose membranes. For western blotting analysis, membranes were incubated with appropriate antibodies at 4°C for overnight followed by incubation with a secondary antibody. Immunoreactive bands were visualized using western blotting Luminal reagent (Santa Cruz Biotechnology) according to the manufacturer’s recommendation.

### Fast Protein Liquid Chromatography (FPLC)

Cellular lysates from MDA-MB-231 cells stably expressing FLAG-ZNF704 were prepared by incubating the cells in lysis buffer containing the protease inhibitor cocktail. Anti-FLAG immunoaffinity columns were prepared using the anti-FLAG M2 affinity gel following the manufacturer’s suggestions. Cell lysates were obtained and applied to an equilibrated FLAG column of 1-ml bed volume to allow for adsorption of the protein complex to the column resin. After binding, the column was washed with cold PBS plus 0.1% Nonidet P-40. FLAG peptide was then applied to the column to elute the FLAG protein complex as described by the vendor. Fractions of the bed volume were collected and concentrated to 0.5 ml using a Millipore Ultrafree centrifugal filter apparatus (3 kDa nominal molecular mass limit), and then applied to an 850 x 20 mm Superose 6 size exclusion column (Amersham Biosciences) that was equilibrated with PBS and calibrated with protein standards (blue dextran, 2000 kDa; thyroglobulin, 669 kDa; ferritin, 440 kDa; aldolase, 158 kDa; ovalbumin, 43 kDa; all from Amersham Biosciences). The column was eluted at a flow rate of 0.5 ml/min and fractions were separately collected. MDA-MB-231 cell nuclear extracts were prepared and dialyzed against buffer (20 mM HEPES, pH 8.0, 10% glycerol, 0.1 mM EDTA, and 300 mM NaCl) (Applygen Technologies Inc.). Approximately 5 mg nuclear proteins were concentrated to 0.5 ml using the Millipore Ultrafree centrifugal filter apparatus.

### GST Pull-down Assay

GST-fused constructs were expressed in BL21 *Escherichia coli*. *In vitro* transcription and translation experiments were done with rabbit reticulocyte lysate (TNT systems, Promega) according to the manufacturer’s recommendation. For GST pull-down assays, about 5 μg of the appropriate GST fusion proteins and 30 μl of glutathione-Sepharose beads were incubated with 5-8 μl of *in vitro* transcribed/translated products in binding buffer (75 mM NaCl, 50 mM HEPES, pH 7.9) at 4°C for 2 h in the presence of the protease inhibitor mixture. The beads were washed 5 times with binding buffer, resuspended in 30 μl of 2 x SDS-PAGE loading buffer, and detected by western blotting.

### RT PCR and Real-time RT PCR (qPCR)

Total cellular RNAs were isolated with TRIzol reagent (Invitrogen) and used for the first strand cDNA synthesis with the Reverse Transcription System (Roche). Quantitation of all gene transcripts was done by qPCR using Power SYBR Green PCR Master Mix and a Roche LightCycler480 sequence detection system with the expression of *GAPDH* as an internal control. The primer pairs used were: *ZNF704*, 5′-GATCAAGCTCAACACAGACTCA-3′ (forward) and 5′-TCTGGGATGGGGAAAGTAGGA-3′ (reverse); *PER2*, 5′-GACATGAGACCAACGAAAACTGC-3′ (forward) and 5′AGGCTAAAGGTATCTGGACTCTG-3′ (reverse); *SIN3A*, 5′-ACCATGCAGTCAGCTACGG-3′ (forward) and 5′-CACCGCTGTTGGGTGATGA-3′ (reverse); *GATA2*, 5′-CAGCAAGGCTCGTTCCTGTT-3′ (forward) and 5′-GGCTTGATGAGTGGTCGGT-3′ (reverse); *CTNNA1*, 5′-GGGGATAAAATTGCGAAGGAGA-3′ (forward) and 5′-GTTGCCTCGCTTCACAGAAGA-3′ (reverse); *FOXO3*, 5′-TCACGCACCAATTCTAACGC-3′ (forward) and 5′-CACGGCTTGCTTACTGAAGG-3′ (reverse); and *GAPDH*, 5′-CTGGGCTACACTGAGCACC-3′ (forward) and 5′-AAGTGGTCGTTGAGGGCAATG-3′ (reverse).

### ChIP-seq

Approximately 5 x 10^7^ cells were used for each ChIP-seq assay. Chromatin DNAs precipitated by polyclonal antibodies against ZNF704 or SIN3A were purified with the Qiagen PCR purification kit. In depth whole-genome DNA sequencing was performed by BGI (Shenzhen, China). The raw sequencing image data were examined by the Illumina analysis pipeline, aligned to the unmasked human reference genome (UCSC GRCh37, hg19) using Bowtie 2, and further analyzed by MACS (Model-based Analysis for ChIP-seq). Genomic distribution of ZNF704 binding sites was analyzed by ChIPseeker, annotated by R package, and compared and visualized(Yu et al, 2015). *De novo* motif screening was performed on sequences ± 100 bp from the centers of ZNF704 or SIN3A binding peaks based on the MEME suite (http://meme-suite.org/). Ontologies analysis was conducted based on the Database for Annotation, Visualization, and Integrated Discovery (DAVID, https://david.ncifcrf.gov/).

### ChIP and Re-ChIP

DNA was purified with the QIAquick PCR Purification Kit. qChIP was performed using the TransStart Top Green qPCR supermix (TransGen Biotech). Re-ChIPs were done essentially the same as primary IP. Bead eluates from the first immunoprecipitation were incubated with 10 mM DTT at 37°C for 30 min and diluted 1:50 in dilution buffer (1% Triton X-100, 2 mM EDTA, 150 mM NaCl, 20 mM Tris-HCl, pH 8.1) followed by re-immunoprecipitation with secondary antibodies. The final elution step was performed using 1% SDS solution in Tris-EDTA buffer, pH 8.1. The sequences of the primers used were: *PER2*, 5′-TTTCCCCAGGCTCTTCTC-3′ (forward) and 5′-GGGAGGCTGTTCTTTGTT-3′ (reverse); *GATA2*, 5′-CAATTACCGACTGTCAATCCCG-3′ (forward) and 5′-TCCTCCAGCCCTCTTCCCT-3′ (reverse); *CTNNA1*, 5′-GGGGACGGGTTAGGTGAA-3′ (forward) and 5′-AGAAGGAGAAGCGGAGGC-3′ (reverse); and *FOXO3*, 5′-TCTGAGCTATCTCCGGTGACTT-3′ (forward) and 5′-CCTTATTCTACGATCCGTGCC-3′ (reverse).

### Time-Series Protein Assay

Time-series protein assay in MDA-MB-231, or U2OS cells was performed as previously described(Balsalobre et al, 2000). Approximate 500,000 cells were plated in 35 mm dishes at 37°C until confluent. Medium was then replaced with serum-free DMEM or L-15 for synchronization of cells for 24 h. The medium was then changed to serum-free DMEM or L-15 with 200 nM dexamethasone (time=0) at 37°C for 2 h and cells were collected at a 4-h interval from 24 to 48 h.

### Lumicycle

Lumicycle analysis of MDA-MB-231- or U2OS-*per2*-luci cells was conducted as previously described(Ramanathan et al, 2012). Briefly, cells were plated in 35-mm dishes at a concentration of 500,000 cells/plate at 37°C until confluent; Medium was replaced with serum-free DMEM or L-15 for synchronization of cells for 24 h and treated with 200 nM dexamethasone at 37°C for 1 h; DMEM containing 1 x Pen/Strep, 200 nm dexamethasone (Sigma), 2% B-27 (Thermo), 1 mM luciferin (Promega), 14.5 mM NaHCO_3_ (Sigma), and 10 mM HEPES (pH 7.2, Thermo) was applied to synchronized cells. Data were collected in a LumiCycle luminometer at 36°C for 5-6 days and analyzed with LumiCycle Analysis software (Actimetrics). Data from the first 24 h cycle was excluded(Liu et al, 2007).

### Cell Viability/Proliferation Assay

For cell proliferation assays, MCF-7 cells were seeded into 96-well plates with an equal volume of medium. On the day of harvest, 10 μl CCK-8 solution was added according to the manufacturer’s protocol (MedChem Express). Plates were incubated at 37°C for 2 h and cell viability was determined by measuring the absorbance at 450 nm wavelength. Each experiment was performed in triplicate and repeated at least three times.

### Colony Formation Assay

MCF-7 cells were maintained in culture media in 6-well plate for 14 days, fixed with 4% paraformaldehyde, and stained with 0.1% crystal violet for colony observation. Each experiment was performed in triplicate and repeated at least three times.

### Cell Invasion Assay

The transwell invasion assay was performed using the transwell chamber (BD biosciences) with a Matrigel-coated filter. Stably-infected MDA-MB-231 cells were cultured in Leibovitz’s L-15 medium with 10% FBS at 37°C without CO_2_. Cells were cultured in serum-free Leibovitz’s L-15 medium for 24 h and harvested. These cells were then washed three times in PBS, resuspended in serum-free media (5 x 10^4^ of cells in 0.5 ml), and plated onto the upper chamber of the transwell. The upper chamber was then transferred into a well containing 0.5 ml of media supplemented with 10% FBS and incubated for 16 h. Cells on the upside were removed using cotton swabs, and the invasive cells on the lower side were fixed, stained with 0.1% crystal violet solution, and counted using light microscopy. Each experiment was performed in triplicate and repeated at last three times.

### *In Vivo* Metastasis

The MDA-MB-231-Luc-D3H2LN cells (Xenogen Corporation) were infected with lentiviruses carrying control shRNA+vector, FLAG-ZNF704, or/and SIN3A shRNA or shCTR, ZNF704 shRNA, or/and PER2 shRNA. These cells were inoculated onto the left abdominal mammary fat pad (3 x 10^6^ cells) or injected into the lateral tail vein (1 x 10^6^ cells) of 6-week-old immunocompromised female SCID beige mice (n=6). Bioluminescent images were obtained with a 15-cm field of view, binning (resolution) factor of 8, 1/f stop, open filter, and an imaging time of 30 s to 2 min. Bioluminescence from relative optical intensity was defined manually. Photon flux was normalized to background which was defined from a relative optical intensity drawn over a mouse not given an injection of luciferin.

### Tissue Specimens

Samples of breast cancer paired with adjacent normal mammary tissues were obtained from surgical specimens from patients with breast cancer. Samples were selected from patients for whom complete information on clinicopathological characteristics was available. Samples were frozen in liquid nitrogen immediately after surgical removal and maintained at −80°C until RNA extraction.

### Statistics

Results were reported as mean ± SD for triplicate experiments unless otherwise noted. SPSS V.19.0, 2-tailed t test, and 1-way ANOVA were used and indicated in figure legends. The correlation coefficients were calculated by SPSS V.19.0. A *P* value less than 0.05 was considered significant. Datasets were downloaded from http://www.ncbi.nlm.nih.gov/geo, and data for Kaplan–Meier survival analysis was from http://kmplot.com/analysis/index.php?p=service&cancer=breast.

### Study approval

All studies were approved by the Ethics Committee of Capital Medical University and informed consent was obtained from all patients. Animal handling and procedures were approved by the Capital Medical University Institutional Animal Care.

## Acknowledgments

This work was supported by grants (81530073 and 81730079 to Y.S.) from the National Natural Science Foundation of China, and a grant (2016YFC1302304 to Y.S.) from the Ministry of Science and Technology of China.

## References

(2012) Comprehensive molecular portraits of human breast tumours. Nature 490: 61–70

Altman BJ, Hsieh A, Gouw AM, Stine ZE, Venkataraman A, Bellovin DI, Diskin SJ, Lu W, Zhang S, Felsher DW (2014) Abstract 2953: Rev-erbα modulates Myc-driven cancer cell growth and altered metabolism. Cancer Research 74: 2953–2953

Bae K, Jin X, Maywood ES, Hastings MH, Reppert SM, Weaver DR (2001) Differential functions of mPer1, mPer2, and mPer3 in the SCN circadian clock. Neuron 30: 525–536

Baggs JE, Price TS, DiTacchio L, Panda S, Fitzgerald GA, Hogenesch JB (2009) Network features of the mammalian circadian clock. PLoS biology 7: e52

Balleine RL, Fejzo MS, Sathasivam P, Basset P, Clarke CL, Byrne JA (2000) The hD52 (TPD52) gene is a candidate target gene for events resulting in increased 8q21 copy number in human breast carcinoma. Genes Chromosomes & Cancer 29: 48–57

Balsalobre A, Brown SA, Marcacci L, ., Tronche F, ., Kellendonk C, ., Reichardt HM, Schütz G, Schibler U, (2000) Resetting of circadian time in peripheral tissues by glucocorticoid signaling. Science 289: 2344–2347

Bass J, Takahashi JS (2010) Circadian Integration of Metabolism and Energetics. Science 330: 1349–1354

Byrne JA, Balleine RL, Schoenberg FM, Mercieca J, Chiew YE, Livnat Y, St HL, Peters GB, Byth K, Karlan BY (2010) Tumor protein D52 (TPD52) is overexpressed and a gene amplification target in ovarian cancer. International Journal of Cancer 117: 1049–1054

Chen C, Sun X, Guo P, Dong XY, Sethi P, Zhou W, Zhou Z, Petros J, Jr FH, Vessella RL (2007) Ubiquitin E3 ligase WWP1 as an oncogenic factor in human prostate cancer. Oncogene 26: 2386–2394

Chen C, Zhou Z, Ross JS, Zhou W, Dong JT (2010) The amplified WWP1 gene is a potential molecular target in breast cancer. International Journal of Cancer 121: 80–87

Chen ST, Choo KB, Hou MF, Yeh KT, Kuo SJ, Chang JG (2005) Deregulated expression of the PER1, PER2 and PER3 genes in breast cancers. Carcinogenesis 26: 1241–1246

Choschzick M, Lassen P, Lebeau A, Marx AH, Terracciano L, Heilenkötter U, Jaenicke F, Bokemeyer C, Izbicki J, Sauter G (2010) Amplification of 8q21 in breast cancer is independent of MYC and associated with poor patient outcome. Modern Pathology 23: 603–610

Ciriello G, Gatza ML, Beck AH, Wilkerson MD, Rhie SK, Pastore A, Zhang H, McLellan M, Yau C, Kandoth C, Bowlby R, Shen H, Hayat S, Fieldhouse R, Lester SC, Tse GM, Factor RE, Collins LC, Allison KH, Chen YY, Jensen K, Johnson NB, Oesterreich S, Mills GB, Cherniack AD, Robertson G, Benz C, Sander C, Laird PW, Hoadley KA, King TA, Perou CM (2015) Comprehensive Molecular Portraits of Invasive Lobular Breast Cancer. Cell 163: 506–519

Courjal F, Theillet C (1997) Comparative genomic hybridization analysis of breast tumors with predetermined profiles of DNA amplification. Cancer Res 57: 4368–4377

Debruyne JP, Noton E, Lambert CM, Maywood ES, Weaver DR, Reppert SM (2006) A clock shock: mouse CLOCK is not required for circadian oscillator function. Neuron 50: 465–477

Dunlap JC (1999) Molecular bases for circadian clocks. Cell 96: 271–290

Fan W, Chen X, Li C, Chen L, Liu P, Chen Z (2014) The analysis of deregulated expression and methylation of the PER2 genes in gliomas. Journal of Cancer Research & Therapeutics 10: 636

Filipski E, King VM, Li X, Granda TG, Mormont MC, Liu X, Claustrat B, Hastings MH, Levi F (2002) Host circadian clock as a control point in tumor progression. J Natl Cancer Inst 94: 690–697

Fu L, Pelicano H, Liu J, Huang P, Lee C (2002) The circadian gene Period2 plays an important role in tumor suppression and DNA damage response in vivo. Cell 111: 41–50

Grimaldi B, Bellet MM, Katada S, Astarita G, Hirayama J, Amin RH, Granneman JG, Piomelli D, Leff T, Sassone-Corsi P (2010) PER2 controls lipid metabolism by direct regulation of PPARgamma. Cell Metab 12: 509–520

Hida A, Koike N, Hirose M, Hattori M, Sakaki Y, Tei H (2000) The human and mouse Period1 genes: five well-conserved E-boxes additively contribute to the enhancement of mPer1 transcription. Genomics 65: 224–233

Hirota T, Lewis WG, Liu AC, Lee JW, Schultz PG, Kay SA (2008) A chemical biology approach reveals period shortening of the mammalian circadian clock by specific inhibition of GSK-3beta. Proceedings of the National Academy of Sciences of the United States of America 105: 20746–20751

Hoffman AE, Chun-Hui Y, Tongzhang Z, Stevens RG, Derek L, Yawei Z, Holford TR, Johnni H, Jennifer P, Yong Z (2010a) CLOCK in breast tumorigenesis: genetic, epigenetic, and transcriptional profiling analyses. Cancer Research 70: 1459–1468

Hoffman AE, Zheng T, Yi CH, Stevens RG, Ba Y, Zhang Y, Leaderer D, Holford T, Hansen J, Zhu Y (2010b) The core circadian gene Cryptochrome 2 influences breast cancer risk, possibly by mediating hormone signaling. Cancer Prevention Research 3: 539

Hwang-Verslues WW, Po-Hao C, Yung-Ming J, Wen-Hung K, Pei-Hsun C, Yi-Cheng C, Tsung-Han H, Fang-Yi S, Liu-Chen L, Serena A (2013) Loss of corepressor PER2 under hypoxia up-regulates OCT1-mediated EMT gene expression and enhances tumor malignancy. Proceedings of the National Academy of Sciences of the United States of America 110: 12331–12336

Iurisci I, Valet F, Giacchetti S, Pierga JY, André F, Cremoux PD, Asselain B, Delaloge S, Thé HD, Spyratos F (2010) Abstract P2-09-26: Circadian Clock Genes in Primary Breast Cancer: Strong Predictors of Pathologic Response on Neoadjuvant Chemotherapy. Cancer Research 70: P2-09-26–P02-09-26

Kochan DZ, Kovalchuk O (2015) Circadian disruption and breast cancer: an epigenetic link? Oncotarget 6: 16866–16882

Kornmann B, Schaad O, Bujard H, Takahashi JS, Schibler U (2007) System-driven and oscillator-dependent circadian transcription in mice with a conditionally active liver clock. PLoS biology 5: e34

Kumamoto T, Seki N, Mataki H, Mizuno K, Kamikawaji K, Samukawa T, Koshizuka K, Goto Y, Inoue H (2016) Regulation of TPD52 by antitumor microRNA-218 suppresses cancer cell migration and invasion in lung squamous cell carcinoma. International Journal of Oncology 49: 1870–1880

Kume K, Zylka MJ, Sriram S, Shearman LP, Weaver DR, Jin X, Maywood ES, Hastings MH, Reppert SM (1999) mCRY1 and mCRY2 are essential components of the negative limb of the circadian clock feedback loop. Cell 98: 193

Liu AC, Welsh DK, Ko CH, Tran HG, Zhang EE, Priest AA, Buhr ED, Singer O, Meeker K, Verma IM, Doyle FJ, Takahashi JS, Kay SA (2007) Intercellular Coupling Confers Robustness against Mutations in the SCN Circadian Clock Network. Cell 129: 605–616

Liu B, Xu K, Jiang Y, Li X (2014) Aberrant expression of Per1, Per2 and Per3 and their prognostic relevance in non-small cell lung cancer. International Journal of Clinical & Experimental Pathology 7: 7863–7871

Luo Y, Wang F, Chen LA, Chen XW, Chen ZJ, Liu PF, li FF, Li CY, Liang W (2012) Deregulated expression of cry1 and cry2 in human gliomas. Asian Pacific journal of cancer prevention : APJCP 13: 5725–5728

Maier B, Wendt S, Vanselow JT, Wallach T, Reischl S, Oehmke S, Schlosser A, Kramer A (2009) A large-scale functional RNAi screen reveals a role for CK2 in the mammalian circadian clock. Genes & development 23: 708–718

Mcdonel P, Demmers J, Tan DW, Watt F, Hendrich BD (2012) Sin3a is essential for the genome integrity and viability of pluripotent cells. Developmental Biology 363: 62–73

Myllykangas S, Bohling T, Knuutila S (2007) Specificity, selection and significance of gene amplifications in cancer. Semin Cancer Biol 17: 42–55

Preitner N, Damiola F, Luis Lopez M, Zakany J, Duboule D, Albrecht U, Schibler U (2002) The Orphan Nuclear Receptor REV-ERBα Controls Circadian Transcription within the Positive Limb of the Mammalian Circadian Oscillator. Cell 110: 251–260

Raeder MB, Birkeland E, Trovik J, Krakstad C, Shehata S, Schumacher S, Zack TI, Krohn A, Werner HM, Moody SE, Wik E, Stefansson IM, Holst F, Oyan AM, Tamayo P, Mesirov JP, Kalland KH, Akslen LA, Simon R, Beroukhim R, Salvesen HB (2013) Integrated genomic analysis of the 8q24 amplification in endometrial cancers identifies ATAD2 as essential to MYC-dependent cancers. PloS one 8: e54873

Ramanathan C, Khan SK, Kathale ND, Xu H, Liu AC (2012) Monitoring cell-autonomous circadian clock rhythms of gene expression using luciferase bioluminescence reporters. Journal of Visualized Experiments Jove 67: e4234–e4234

Reppert SM, Weaver DR (2002) Coordination of circadian timing in mammals. Nature 418: 935–941

Sandrelli F, Cappellozza S, Benna C, Saviane A, Mastella A, Mazzotta GM, Moreau S, Pegoraro M, Piccin A, Zordan MA, Cappellozza L, Kyriacou CP, Costa R (2007) Phenotypic effects induced by knock-down of the period clock gene in Bombyx mori. Genet Res 89: 73–84

Schernhammer ES, Laden F, Speizer FE, Willett WC, Hunter DJ, Kawachi I, Colditz GA (2001) Rotating night shifts and risk of breast cancer in women participating in the nurses’ health study. J Natl Cancer Inst 93: 1563–1568

Shan L, Zhou X, Liu X, Wang Y, Su D, Hou Y, Yu N, Yang C, Liu B, Gao J, Duan Y, Yang J, Li W, Liang J, Sun L, Chen K, Xuan C, Shi L, Wang Y, Shang Y (2016) FOXK2 Elicits Massive Transcription Repression and Suppresses the Hypoxic Response and Breast Cancer Carcinogenesis. Cancer Cell 30: 708–722

Shih MC, Yeh K-T, Tang K-P, Chen J-C, Chang J-G (2010) Promoter methylation in circadian genes of endometrial cancers detected by methylation-specific PCR. Mol Carcinog 45: 732–740

Si W, Huang W, Zheng Y, Yang Y, Liu X, Shan L, Zhou X, Wang Y, Su D, Gao J, Yan R, Han X, Li W, He L, Shi L, Xuan C, Liang J, Sun L, Wang Y, Shang Y (2015) Dysfunction of the Reciprocal Feedback Loop between GATA3- and ZEB2-Nucleated Repression Programs Contributes to Breast Cancer Metastasis. Cancer Cell 27: 822–836

Streubel G, Fitzpatrick DJ, Oliviero G, Scelfo A, Moran B, Das S, Munawar N, Watson A, Wynne K, Negri GL (2017) Fam60a defines a variant Sin3a-Hdac complex in embryonic stem cells required for self-renewal. Embo Journal 36: 2216

Sun CM, Huang SF, Zeng JM, Liu DB, Xiao Q, Tian WJ, Zhu XD, Huang ZG, Feng WL (2010) Per2 inhibits k562 leukemia cell growth in vitro and in vivo through cell cycle arrest and apoptosis induction. Pathology oncology research : POR 16: 403–411

Takahashi JS (2016) Transcriptional architecture of the mammalian circadian clock. Nature Reviews Genetics 18: 164

Taniguchi H, Fernández AF, Setién F, Ropero S, Ballestar E, Villanueva A, Yamamoto H, Imai K, Shinomura Y, Esteller M (2009) Epigenetic Inactivation of the Circadian Clock Gene BMAL1 in Hematologic Malignancies. Cancer Research 69: 8447–8454

Tirkkonen M, Tanner M, Karhu R, Kallioniemi A, Isola J, Kallioniemi OP (1998) Molecular cytogenetics of primary breast cancer by CGH. Genes Chromosomes Cancer 21: 177–184

Viswanathan AN, Hankinson SE, Schernhammer ES (2007) Night shift work and the risk of endometrial cancer. Cancer Res 67: 10618–10622

Wang Q, Ao Y, Yang K, Tang H, Chen D (2016) Circadian clock gene Per2 plays an important role in cell proliferation, apoptosis and cell cycle progression in human oral squamous cell carcinoma. Oncol Rep 35: 3387–3394

Wang Y, Shang Y (2013) Epigenetic control of epithelial-to-mesenchymal transition and cancer metastasis. Exp Cell Res 319: 160–169

Wang Y, Zhang H, Chen Y, Sun Y, Yang F, Yu W, Liang J, Sun L, Yang X, Shi L, Li R, Li Y, Zhang Y, Li Q, Yi X, Shang Y (2009) LSD1 is a subunit of the NuRD complex and targets the metastasis programs in breast cancer. Cell 138: 660–672

Winter SL, Bosnoyan-Collins L, Pinnaduwage D, Andrulis IL (2007) Expression of the circadian clock genes Per1 and Per2 in sporadic and familial breast tumors. Neoplasia 9: 797–800

Xiong H, Yang Y, Yang K, Zhao D, Tang H, Ran X (2018) Loss of the clock gene PER2 is associated with cancer development and altered expression of important tumor-related genes in oral cancer. International Journal of Oncology 52: 279

Yan R, He L, Li Z, Han X, Liang J, Si W, Chen Z, Li L, Xie G, Li W, Wang P, Lei L, Zhang H, Pei F, Cao D, Sun L, Shang Y (2015) SCF(JFK) is a bona fide E3 ligase for ING4 and a potent promoter of the angiogenesis and metastasis of breast cancer. Genes Dev 29: 672–685

Yang X, Wood PA, Oh EY, Du-Quiton J, Ansell CM, Hrushesky WJ (2009) Down regulation of circadian clock gene Period 2 accelerates breast cancer growth by altering its daily growth rhythm. Breast Cancer Res Treat 117: 423–431

Yeh CM, Shay J, Zeng TC, Chou JL, Huang TH, Lai HC, Chan MW (2014) Epigenetic silencing of ARNTL, a circadian gene and potential tumor suppressor in ovarian cancer. International Journal of Oncology 45: 2101–2107

Yu EA, Weaver DR (2011) Disrupting the circadian clock: gene-specific effects on aging, cancer, and other phenotypes. Aging 3: 479–493

Yu G, Wang LG, He QY (2015) ChIPseeker: an R/Bioconductor package for ChIP peak annotation, comparison and visualization. Bioinformatics (Oxford, England) 31: 2382–2383

Yu H, Meng X, Wu J, Pan C, Ying X, Zhou Y, Liu R, Huang W (2013) Cryptochrome 1 overexpression correlates with tumor progression and poor prognosis in patients with colorectal cancer. Plos One 8: e61679

Zhang Y, Zhang D, Li Q, Liang J, Sun L, Yi X, Chen Z, Yan R, Xie G, Li W, Liu S, Xu B, Li L, Yang J, He L, Shang Y (2016) Nucleation of DNA repair factors by FOXA1 links DNA demethylation to transcriptional pioneering. Nat Genet 48: 1003–1013

Zhao H, Zeng ZL, Yang J, Jin Y, Zou Q-F (2014) Prognostic relevance of Period1 (Per1) and Period2 (Per2) expression in human gastric cancer. International Journal of Clinical & Experimental Pathology 7: 619–630

